# It’s complicated: how sex, family, and season affect growth in a sexually size-dimorphic spider

**DOI:** 10.64898/2025.12.04.692355

**Authors:** Tim Prezelj, Paul Vincent Debes, Rok Golobinek, Simona Kralj-Fišer

## Abstract

Female-biased sexual size dimorphism (SSD), where females are larger than males, is widespread. While ultimate explanations for SSD are well established, the proximate developmental mechanisms remain poorly understood. Studying sex-specific growth trajectories to identify common vs. sex-specific growth periods is therefore key to uncovering when and how SSD emerges. Theory predicts that female-biased SSD may arise if females hatch larger, grow more rapidly, grow longer, or combine these pathways.

We studied sex-specific growth trajectories in the African hermit spider, *Nephilingis cruentata*, where adult females are on average 75-times heavier and take 2.3-times longer until adulthood than males. We tracked growth trajectories of carapace width for 916 individuals. After initial growth trajectories common to both sexes, trajectories diverged after ∼80 days, coinciding with the onset of the male subadult stage. At this point, male growth decelerated and terminated at sexual maturity, whereas female growth accelerated until their subadult stage at ∼200 days. Thereafter, female growth rate also decelerated and eventually terminated at sexual maturity. Thus, females attain their larger size through both an extended growth period and a more rapid growth rate, which initiates in mid-development. Whereas we detected seasonal effects on growth that were similar in both sexes, family effects showed strong sex-specific signatures. Our results pinpoint a key developmental window in which male and female growth trajectories begin to diverge, providing a clear target for detailed investigation of the mechanisms underlying SSD. At the same time, the complexity of these patterns will likely hinder mechanistic studies of SSD development.

## INTRODUCTION

Sexual dimorphism—the phenotypic difference between males and females—is widespread across animals and often manifests in coloration, behaviour, morphology, or body size (Cunningham 1900; Hedrick & Temeles 1989; Shine 1989; Fairbairn et al. 2007). Among these traits, sexual size dimorphism (SSD) has attracted particular attention. While males are typically the larger sex in most birds and mammals (e.g., great bustard, elephant seals), females are generally larger in many ectothermic taxa, including most spiders (Fairbairn 1997; Lindenfors et al. 2002; Kuntner & Coddington 2020).

Spiders exhibit the most pronounced female-biased sexual size dimorphism (SSD) among terrestrial animals, establishing them as an excellent model system for investigating SSD evolution. Comparative surveys indicate that approximately 85% of species display female-biased SSD (Kuntner & Coddington 2020). In certain lineages, this dimorphism reaches extraordinary levels, with females exceeding males by up to 1000-fold in body mass (Kuntner & Coddington 2020). As another advantage to the study of SSD, spiders also exhibit discrete and determinate growth, ceasing to grow upon reaching sexual maturity (Foelix 2010).

Evolutionary explanations for female-biased SSD generally emphasize fecundity selection, whereby larger females achieve higher reproductive output (Head 1995; Prenter et al. 1999; but see Higgins et al. 2011). Several hypotheses rationalize why males remain small, such as early maturation through scramble competition (Christenson & Goist 1979; Kasumovic & Andrade 2009; Danielson-François et al. 2012), reduced risk of sexual cannibalism (Elgar et al. 2000; Kralj-Fišer et al. 2016), and enhanced agility during mate searching (Moya-Laraño et al. 2002; Corcobado et al. 2010; but see Quiñones-Lebrón et al. 2019). Empirical tests of these hypotheses, however, have yielded mixed results.

While SSD has been broadly studied from evolutionary and ecological perspectives, i.e., its ultimate explanations, its proximate developmental mechanisms remain less well understood. SSD can arise through several developmental mechanisms, i.e., the larger sex may hatch larger, grow more rapidly, develop for a longer period, or reach a larger size through a combination of these mechanisms (Roff 1993; Higgins & Rankin 1996; Blanckenhorn et al. 2007). Sex differences in egg or hatchling size are comparatively rare (Stillwell & Davidowitz 2010; Teder 2014, but see Budrienė et al. 2013). In arthropods, the larger sex typically prolongs development (e.g., Higgins 2000; Teder 2014). In other cases, the larger sex grows more rapidly during an equivalent developmental period (Vendl et al. 2018) or combines a prolonged growth duration with accelerated growth (Sõber et al. 2019; Agnarsson et al. 2024). Growth is further shaped by environmental variation, including food, temperature, and photoperiod, with developmental plasticity often differing between sexes (Leimar 1996; Li & Jackson 1996; Nylin & Gotthard 1998; Fernández-Montraveta & Moya-Laraño 2007; Neumann et al. 2017; Andrade 2019; Lissowsky et al. 2021).

Whereas in most spiders exhibiting female-biased SSD, females extend their growth period and, in some species, also achieve faster growth rates (Higgins 2000; Uhl et al. 2004; Kralj-Fišer et al. 2014; Zhang et al. 2021; Agnarsson et al. 2024), the developmental stage or timing of sex-specific divergence in body size remains poorly resolved. Two well-studied cases illustrate contrasting patterns. In the swift crab spider (*Mecaphesa celer*), males and females grow similarly during early instars, but from the fifth instar onward females both accelerate and prolong growth, ultimately attaining much larger adult sizes (Chelini et al. 2019). In contrast, in the yellow garden spider (*Argiope aurantia*), females already outpace males during early juvenile stages, indicating an earlier onset of sex-specific growth trajectories (Inkpen & Foellmer 2010). Together, these examples demonstrate that SSD can arise, depending on the species, through either late divergence between sexes in growth or early developmental growth rate differences, thereby highlighting the need for comprehensive ontogenetic studies across spider lineages.

Here we investigate sex-specific growth trajectories in the African hermit spider (*Nephilingis cruentata*, Fabricius, 1775), which exhibits extreme SSD (**Figure 1**). Specifically, females are ∼75 times heavier and develop 2.3 times longer than males (Šet et al. 2021; Kralj-Fišer et al. 2023). While it is well established that females develop longer and attain larger adult body mass than males (Šet et al. 2021; Kralj-Fišer et al. 2023), longitudinal growth trajectories from hatching to sexual maturity (i.e., adulthood) remain unknown for either sex. Thus, it is unclear whether females also grow more rapidly than males, and at what developmental stage their growth trajectories may begin to diverge. However, growth rates of some spiders vary with season (Higgins 2000; Lissowsky et al. 2021) and this complexity in studying growth has previously been observed for adult body mass variation in our study species (Kralj-Fišer et al. 2023).To study growth differences between sexes and across seasons in *N. cruentata*, we recorded body size longitudinally for both sexes from hatching to sexual maturity across a three-year period, using carapace width as a linear proxy for body size (Fernández-Montraveta & Marugán-Lobón 2017).

**Figure 1.**
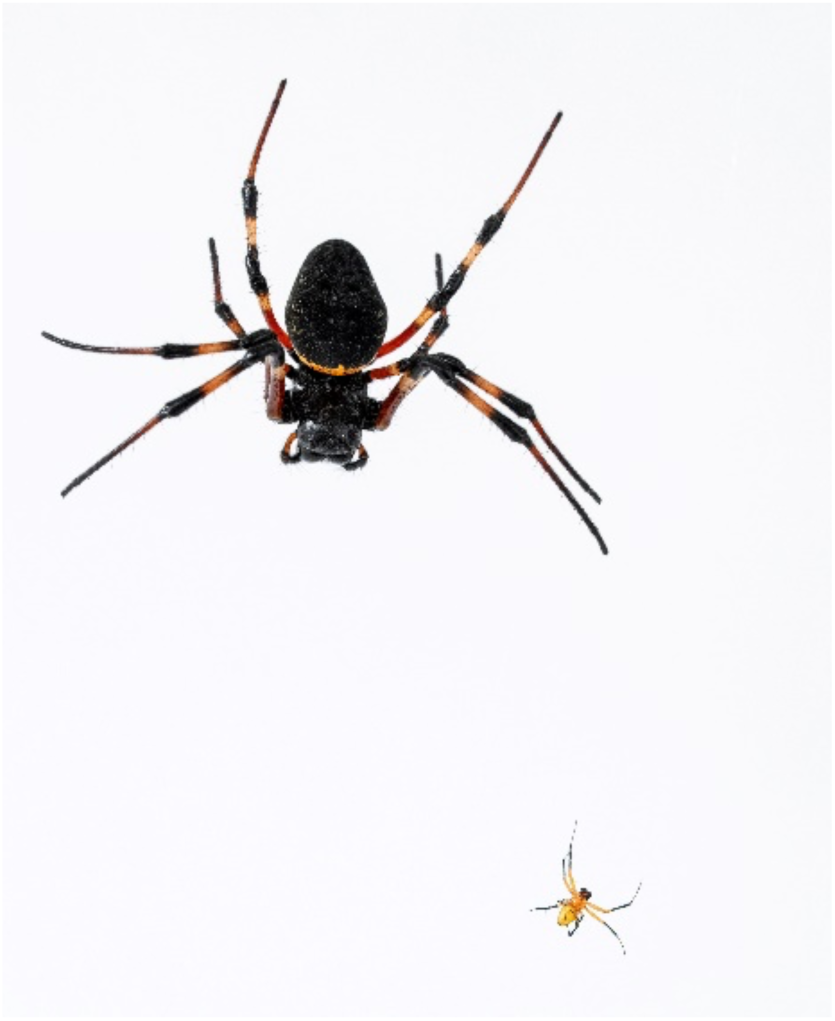
Female (large) and male (small) African hermit spider (*Nephilingis cruentata*). Photo by Manca Juvan.

Because we already know that in our species females take longer to reach adulthood than males, we tested the hypotheses that females, relative to males, (i) achieve larger adult size by also growing more rapidly, and (ii) that seasonal growth effects on adult body mass extend to carapace-width growth. Importantly, we explored when during development females and males diverge in their growth trajectories and size. Our study therefore allows reliably pinpoint when and how male and female growth trajectories diverge, which is essential for understanding the proximate developmental basis of SSD and also provides a foundation for future studies linking these processes to molecular and genetic mechanisms.

## MATERIALS AND METHODS

### Study system and rearing

We established laboratory colonies of the study species, *N. cruentata* in 2015 from gravid females collected in iSimangaliso Wetland Park and Ndumo Reserve, South Africa (permit OP 552/2015), with additional spiders added in 2018 (permit OP 3031/2020). Spiders were reared under standardized conditions (25 ± 2 °C, 45 ± 5% humidity, 12:12 h light–dark cycle with LED and natural light). To maintain the colony, we paired randomly selected males and females (avoiding siblings), and reared offspring from the first egg sac per female (see details in Kralj-Fišer et al. 2023). We monitored egg sacs twice weekly and isolated spiderlings two weeks post-hatching. Juveniles were fed *Drosophila* until the 4^th^ moult and thereafter blowflies (*Lucilia sericata*).

### Data collection

To obtain carapace width estimates, we photographed spiders dorsally with a Canon EOS 7D + 50 mm macro lens under fixed settings. We calculated carapace width (mm) from calibrated pixel measurements using the equation: 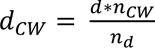 , where *d_CW_* is carapace width; *d* is the absolute distance; *n_CW_* is carapace width in pixels; and *n_d_* is pixels representing the chosen absolute distance Quiñones-Lebrón et al. 2021).

In total, we measured 916 spiders (n females = 256, n males = 658). Although we aimed to measure carapace width of each individual repeatedly, this was not possible for all individuals due to death or logistical constraints. As a result, we measured 119 females and 145 males only once. We measured some individuals up to four times covering the developmental stages of juveniles, subadults and adults. *Juveniles* comprise all earlier instars prior to the subadult stage. *Subadults* are individuals at one moult prior to sexual maturity; subadult males can be recognized by enlarged pedipalps, whereas subadult females exhibit a developing, sclerotized epigynal plate beneath the exoskeleton. *Adult* individuals are sexually mature and no longer moult or grow. We provide a full breakdown of sample sizes by sex and life stage in Table S1. In addition to carapace width, we recorded adult body mass within two days of reaching sexual maturity using a precision scale (KERN ABT 100-5NM, d = 0.00001 g).

### Data Analysis

To test for sex differences in growth trajectories for carapace width, we employed a longitudinal general linear mixed model. We modelled the response of ln-transformed carapace width as a function of sex (factor; male or female), age (continuous variable; days after hatch), the age-by-sex interactions, hatching day (continuous variable; day of the year), and the hatching day-by-sex interactions. We fitted hatching day because season affects adult body mass in this species (Kralj-Fišer et al. 2023) and may thus also affect carapace width growth. To capture non-linear growth curves, we fitted age affects as fourth-order polynomials. To capture non-linear season effects, we followed Kralj-Fišer et al. (2023) and fitted hatching day as third-order polynomials. To account for correlations among carapace width measurements due to individual permanent environmental effects, we included *individual* (*ind*) as random intercept effects. To account for correlations among carapace width measurements due to relatedness among spiders, we fitted *family* (*fam*) as random intercepts and as first-order polynomial effects for age (i.e., linear slopes). Assuming the first order polynomials capture most of the variation in growth rates and higher order polynomials mostly curvature, we thus allowed the different families to have both different average size and different growth rates. We allowed the variance for the permanent environmental *individual* effects, and for the model *residuals* (*err*) to vary between sexes. We also fitted a covariance between sex-specific *family* effects but not between sex-specific *individual* or *residual* effects because these effects were not estimable. In addition, we fitted covariances between the *family* intercepts and slopes, resulting in a 4 x 4 covariance matrix for sex-specific *family* intercepts and age slopes, 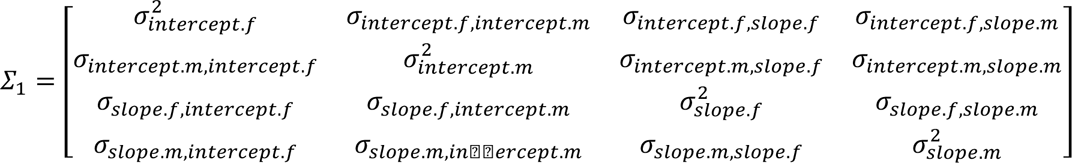 where subscript *f* indicates female and *m* male effects. For sex-specific *individual* and *residual* effects, we fitted simpler 2 x 2 covariance matrices, 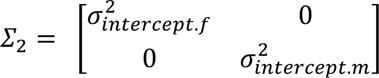. Accordingly, and assuming multivariate normal distributions for all random effects, we modelled *fam* ∼MVN (0, *∑*_1_ ⊗ *F*), *ind* ∼MVN (0, *∑*_2_ ⊗ *I*), and *err* ∼MVN (0, *∑*_2_ ⊗ *E*), where *F*, *I*, and *E* are the respective (sex-specific) identity matrices. The fitted model including a model intercept was, with random effects in italic: intercept + sex + age + sex:age + hatching day + sex:hatching day + sex:*ind* + sex:*fam* + sex:*fam:age* + sex:*err* (*1*).

We also estimated how well size measurements for adult carapace width agreed with those for adult body mass – an often-debated topic. We did so by fitting a multivariate mixed model to the log (ln) of carapace width and the log (ln) of body mass for a data subset of the data, namely for only adult records. This subset consists of 192 female records and 562 male records. The mixed model was fitted with trait-specific fixed effects for sex, hatching day, and the sex-by-hatching day interaction. No age effects were fitted because we only used adult records which do not require modelling temporal effects, i.e., growth. We fitted *fam* as random effects, and allowed the among-family variance to differ among sexes and traits and have covariance between sexes and traits via a 4 x 4 covariance matrix 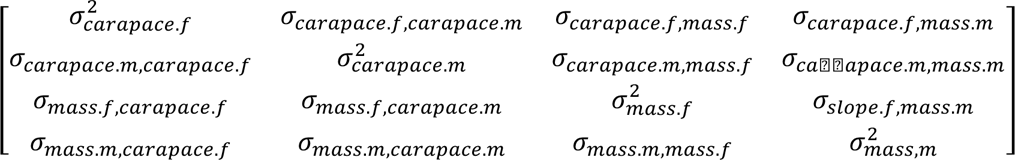model residuals (*err*) were also allowed to differ among sexes and traits and to have a covariance between traits, but not between sexes, because these effects were not estimable, via a 4 x 4 covariance matrix 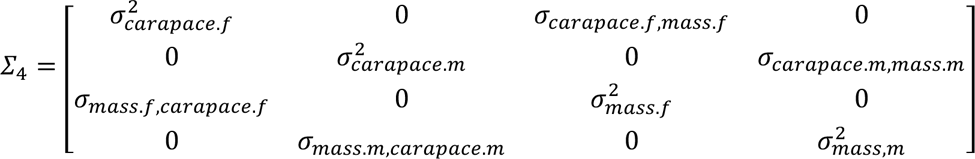 assumed multivariate normal distributions for all random effects. The fitted model including model intercepts per trait (trait) was thus, with random effects in italic: trait + trait:sex + trait:hatching day + trait:sex:hatching day + *trait:sex:fam* + *trait:sex:err* (*2*).

From the fitted longitudinal mixed model (*1*), we predicted average effects, their standard errors, and standard errors of differences via the *asreml-R* “predict” function. In detail, we predicted sex-specific growth trajectories (age affects) for carapace width, the difference between the sexes, and sex-specific seasonal hatching day effects on adult carapace width. We also predicted sex-specific familial growth trajectories (age affects) for carapace width, but only for families with a minimum of three females and three males covering data until the adult stage. To predict sex-specific growth trajectories, and the standard error of the sex difference, we fixed the hatching day effect at the overall average value across sexes (day 177, corresponding to 26^th^ or 27^th^ of June across the experimental period between 2020 and 2023). To predict season effects on adult carapace width, we fixed the age effect to the average number of days (d) it took to reach the adult stage for each sex (females: 221.1 d, males: 94.3 d). We calculated approximate confidence intervals as estimates ± two times the standard error.

From the fitted multivariate model (*2*), we extracted correlations, their standard errors, the difference between the trait-specific between-sex correlations, and the standard errors of this difference via the *asreml-R* “vpredict” function (which implements the delta method). We calculated correlations for family and residual effects as the ratio of the respective covariance to the square root of the respective variance product and the phenotypic correlation as the ratio of the covariance sum (for family and residual effects) to the variance product of the variance sums (for family and residual effects).

## RESULTS

The estimated female growth trajectories showed that females grew more rapidly after hatching and before reaching sexual maturity than in mid-development (**Figure 2a**). The estimated male growth trajectories showed that males followed a simpler function than females. Males were initially growing with the highest growth rate that slowed down evenly until reaching final size at sexual maturity (**Figure 2b**).

**Figure 2.**
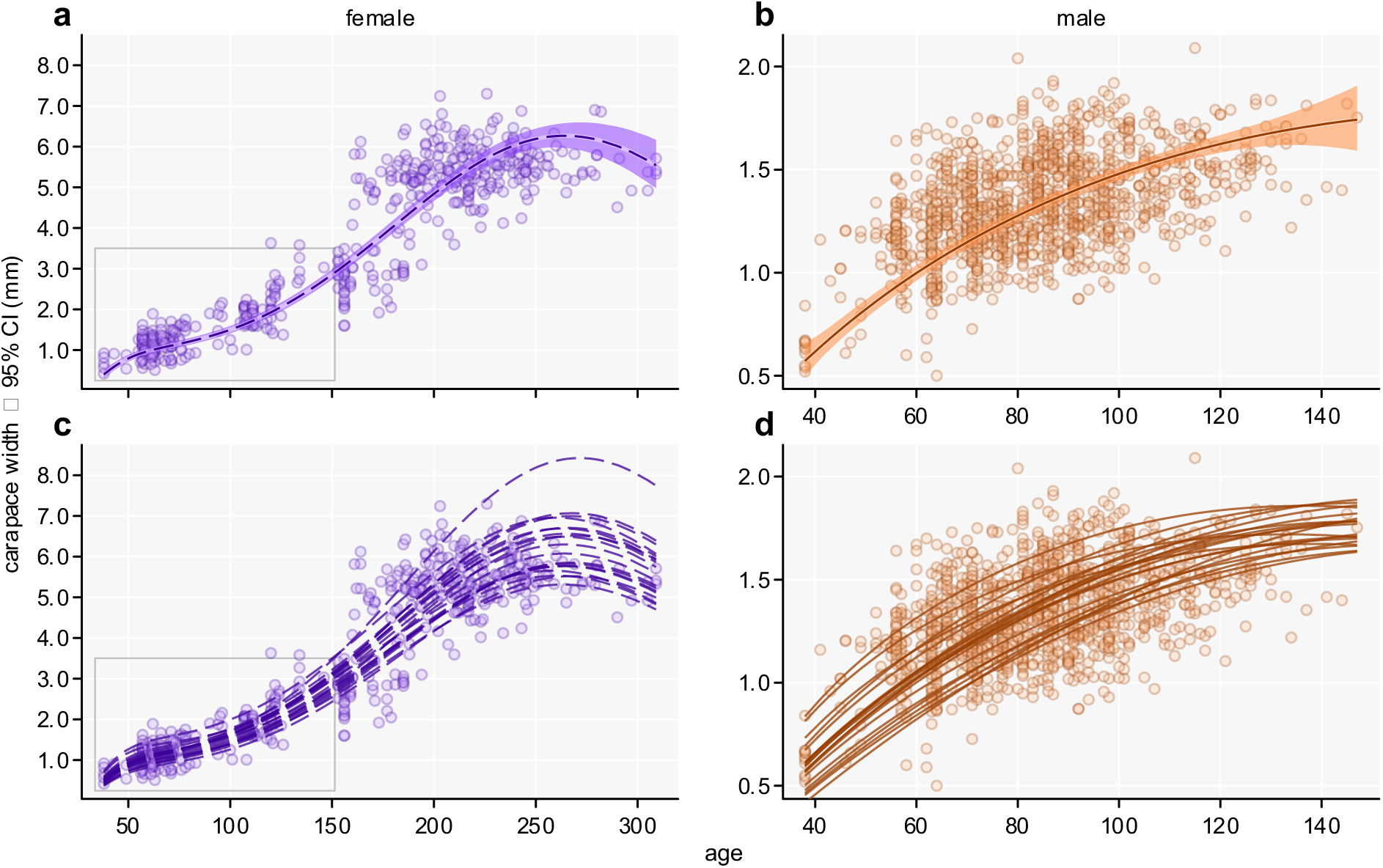
Mixed-model predicted sex-specific growth trajectories of *Nephilingis cruentata* from hatch until adulthood. Shown are the average trajectories with 95% confidence bands (**a**, **b**) and the family-specific trajectories for families with at least three individuals reaching the adult stage per sex (**c**, **d**). The points show data records and the grey box in the female panels (**a**, **c**) shows the extent of the male panels (**b**, **d**).

We detected evidence for different growth trajectories between the sexes. Specifically, most of the fitted fourth-order polynomials for age, and their interactions with sex, were significantly different from zero (**Table 1**). A closer visual inspection of the sex-specific growth trajectories for the period that is shared between sexes indicated that females did not grow more rapidly than males for about two-third of the male growing period (**Figure 3**). A similar growth was also indicated by comparing the age-specific size between sexes; the average carapace width was not statistically different between sexes between ages of about 45 and 105 days (**Figure 4**). However, because different growth rates must predate different sizes, the entire period with no statistically significant differences in size cannot be taken as an indication for no statistically significant difference in growth rates. From the average growth trajectories, a divergence between female and male growth can be visually identified around the age of 90 days (**Figure 2b**). Taking an additional account of the large among-family variation for female and male growth trajectories (**Figure 3c, d**), an acceleration of female relative to male growth, and a deceleration of male relative to female growth, may be expected between some families as early as the age of about 70 days and as late as about 110 days. This large temporal among family variation in the onset of either accelerating growth (females) or decelerating growth (males) can be expected to complicate studies on the underlying mechanism of growth and thus size divergence between sexes.

**Figure 3.**
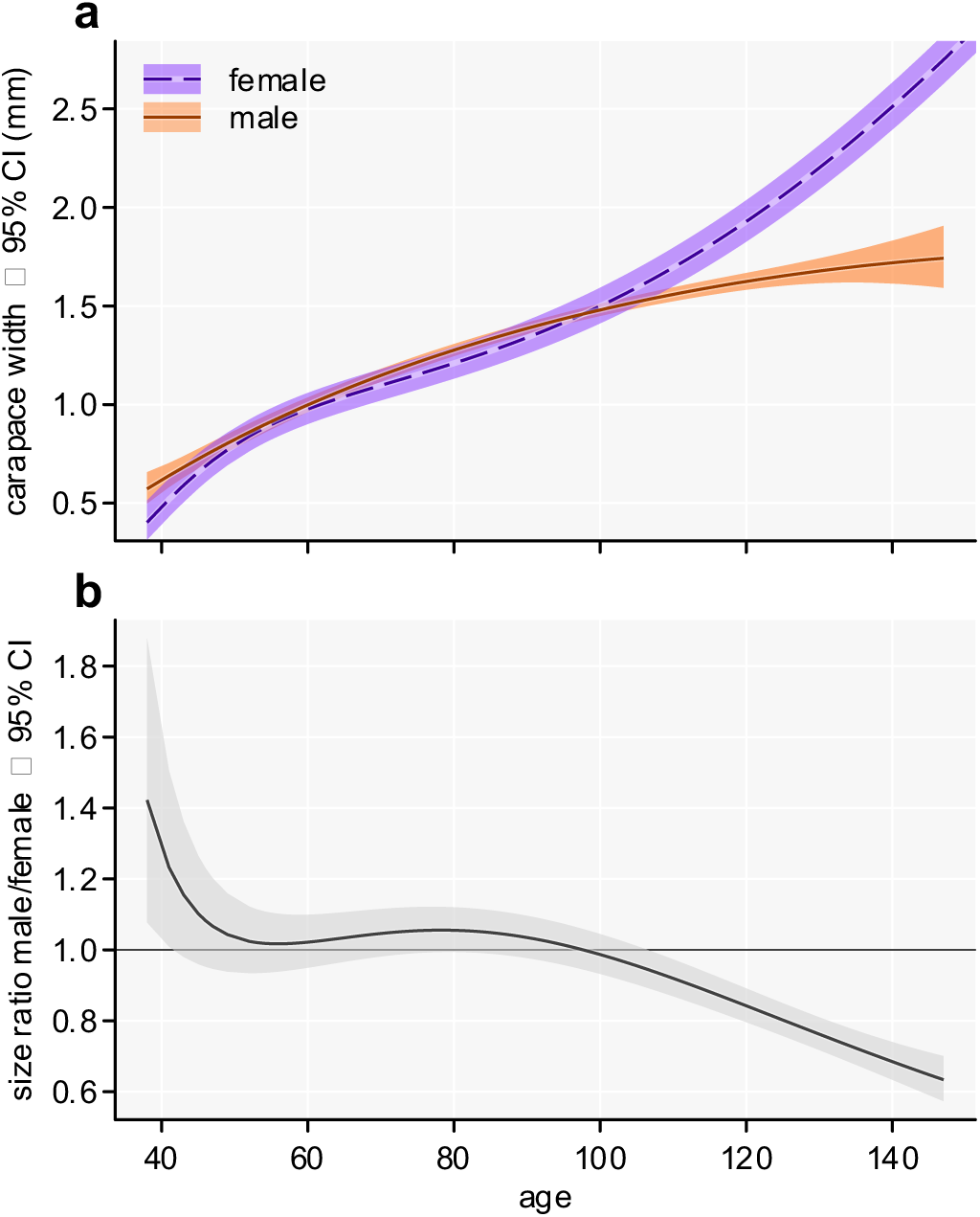
Mixed-model predicted sex-specific growth trajectories for carapace width of *Nephilingis cruentata* but only for the male extent (**a**) and the corresponding size ratio (**b**). Shown are the average trajectories with 95% confidence bands of the estimates (**a**) and the ratio of male over female carapace width with 95% confidence bands of the ratio (**b**), where the horizontal line at one reflects equal size.

**Figure 4.**
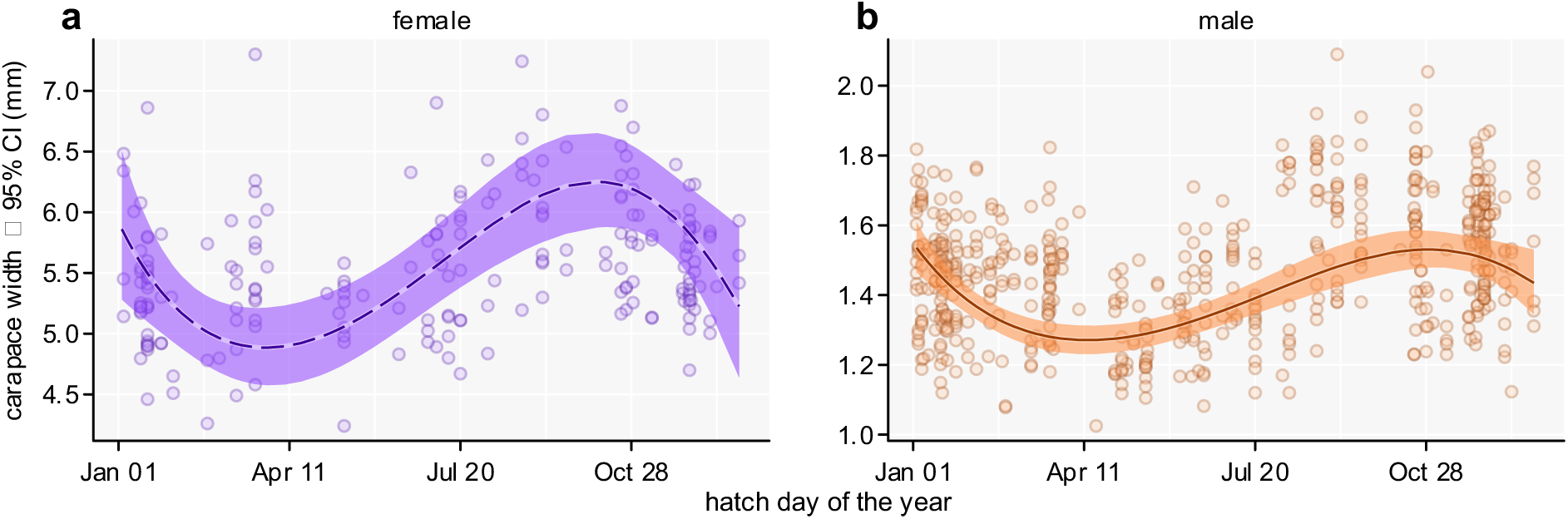
Mixed-model predicted sex-specific adult carapace width of *Nephilingis cruentata* across the day of year when hatched (converted to date for depiction). Shown are the average seasonal curves with 95% confidence bands and points show data records of adult individuals. Curves are predicted for the average adult age per sex (221.1 days for females, 94.3 days for males).

**Table 1.** Model term *F*-test results.

| Term | DF | DDF | F | P |
| --- | --- | --- | --- | --- |
| Sex | 1 | 1641.0 | 19.6 | <0.001 |
| age(1) | 1 | 1641.0 | 2463.0 | <0.001 |
| age(2) | 1 | 897.0 | 0.8 | 0.367 |
| age(3) | 1 | 1178.7 | 18.5 | <0.001 |
| age(4) | 1 | 1028.6 | 83.0 | <0.001 |
| age(1):Sex | 1 | 1234.1 | 114.8 | <0.001 |
| age(2):Sex | 1 | 1080.5 | 67.6 | <0.001 |
| age(3):Sex | 1 | 1250.7 | 25.5 | <0.001 |
| age(4):Sex | 1 | 1172.0 | 5.3 | 0.021 |
| hatch day(1) | 1 | 1641.0 | 17.1 | <0.001 |
| hatch day(2) | 1 | 1641.0 | 21.3 | <0.001 |
| hatch day(3) | 1 | 1641.0 | 33.6 | <0.001 |
| hatch day(1):Sex | 1 | 1641.0 | 0.7 | 0.402 |
| hatch day(2):Sex | 1 | 1641.0 | 1.6 | 0.203 |
| hatch day(3):Sex | 1 | 1641.0 | 1.2 | 0.281 |

Furthermore, the correlation pattern between sexes for the family effects showed an inconsistent pattern between sexes, which complicates meaningful sex contrast within families at specific ages. Specifically, in females we detected a negative correlation -0.74 ± 0.11 (estimate ± standard error) between size intercepts and first-order age slopes whose confidence does not include zero (-0.96 to -0.53), but in males a positive correlation 0.25 ± 0.22 whose confidence interval includes zero (-0.20 to 0.70) and does not overlap with that of females.

Season had a statistically significant effect on carapace width of each sex and these effects, modelled as size proportional effects, did not differ significantly between sexes (**Table 2**). A visual inspection of the seasonal curves for the average adult age per sex indicated that individuals express the smallest average adult body size when hatching around the day of the year 80 (females; second half of March) and 100 (males; first half of April) and the largest average body size when hatching around the day of the year 275 (females; beginning of October) and 290 (males; mid-October).

**Table 2.**
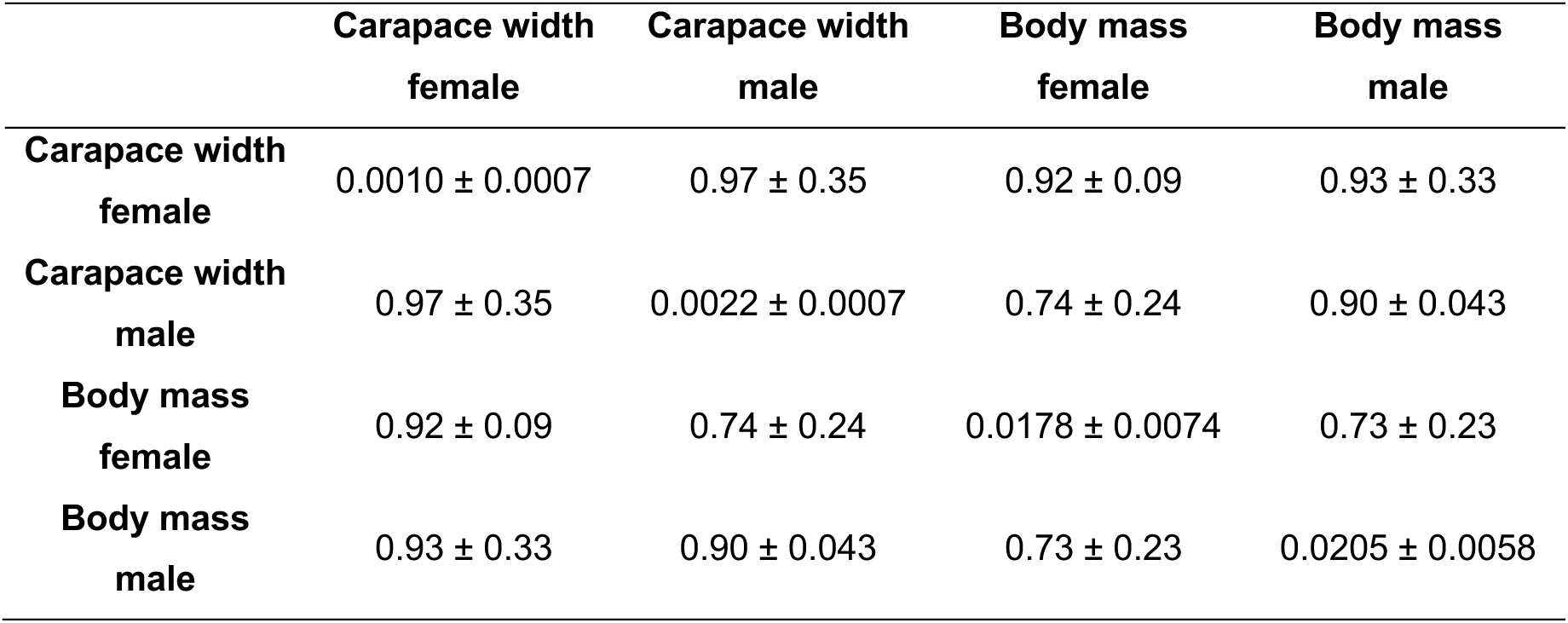
Sex- and trait-specific family effect variance estimates, and between-trait and between-sex correlation estimates, each with standard error, for adult body size based on adult carapace width and adult body mass.

When we modelled adult carapace width and adult body mass in a common multivariate mixed model that took an account of sex- and trait-specific seasonal effect, we found that the partial correlation estimates at both the family and the individual (residual) levels were relatively high (**Table 2**). The between-trait correlation for family effects was with 0.92 and 0.90 for females and males, respectively, similarly high for both sexes. However, the between-sex correlation for family effects was with 0.97 extremely high for carapace width and with 0.73 somewhat lower for body mass, hinting that family-specific sex differences may be more important for body mass than carapace width. However, the difference of the trait-specific between-sex correlations did not significantly differ from zero (difference of the correlations; 95% confidence interval: 0.21; -0.32 to 0.74). The residual correlations between traits were 0.88 ± 0.02 (estimate ± standard error) for female and 0.82 ± 0.02 for males also relatively high and similar between sexes. The calculated sex-specific phenotypic correlation estimates between carapace width and body mass were 0.87 ± 0.02 for females and with 0.83 ± 0.02 for males relatively high and similar between sexes.

## DISCUSSION

Our results provide partial support for the first hypothesis that females attain larger adult size by growing not only for longer than males but also more rapidly. This more rapid female growth occurred during the female mid-phase of development, whereas early growth does not differ between the sexes. Divergence in growth trajectories emerged only around 80–90 days of age, when males reached the subadult stage. We confirm the second hypothesis that seasonal effects on body mass extend to carapace width.

### Sex-specific growth trajectories

Our study in *N. cruentata* shows that male and female growth trajectories were similar during early development but gradually diverged over time. Consistent with this, we detected no statistically significant differences in mean carapace width between sexes from approximately 45 to 105 days of age. Divergence becomes apparent after about 80 days, coinciding with the onset of the subadult stage in males (morphologically observed by enlarged pedipalps). At this point, males begin to decelerate and soon terminate growth, whereas females accelerate growth and continue to grow until their own subadult stage at around 200 days, after which growth also slows and ceases at sexual maturity. In our dataset, the mean age at adulthood was ∼221 days for females and ∼94 days for males. These patterns indicate that females are not growing more rapidly than males after hatching; rather, both sexes share a similar early ontogenetic trajectory. Female-biased SSD thus arises because females extend their growth over a much longer period and exhibit an additional phase of accelerated growth in mid-development, at a time when males are already approaching the end of their growth and development.

Comparative studies suggest that female-biased SSD in spiders can arise through diverse and overlapping developmental mechanisms, some of which are similar to our results. In the pond wolf spider, *Pardosa pseudoannulata*, females undergo an extra moult and grow faster in later instars, with divergence linked to early gonadal differentiation (Zhang et al. 2021). Cupboard spider, *Steatoda grossa*, females also moult more often and, under high food, grow more rapidly than males (Harvey 2022). In the swift crab spider, *Mecaphesa celer*, the two sexes grow similarly at first, but from the fifth instar females accelerate growth relative to males and ultimately reach the larger size (Chelini et al. 2019), which appears similar to what we found in our study species. By contrast, in the yellow garden spider, *Argiope aurantia*, sex-specific differences appear very early in juvenile development, with females already outpacing males early on (Inkpen & Foellmer 2010). Finally, in the golden silk orb-weaver, *Trichonephila clavipes*, females add instars and grow more rapidly than males, producing extreme SSD and pronounced developmental asynchrony (Agnarsson et al. 2024). Together, these examples demonstrate that SSD can originate from different combinations of extended development, increased growth rate, and timing of divergence, highlighting the existence of multiple developmental pathways across spider lineages. Most of all, these differences emphasize the necessity to identify species-specific mechanisms and growth divergence periods that enable more detailed studies on proximate developmental mechanisms underlying SSD.

The timing of sex-specific divergence in growth is central to understanding the proximate mechanisms of SSD. In *P. pseudoannulata* and *A. aurantia*, divergence arises early during juvenile instars, whereas in *N. cruentata* as identified here, *M. celer*, and *S. grossa* it becomes apparent only later, coinciding with the appearance of male secondary sexual traits, such as enlarged pedipalps at the subadult stage. The coincidence of these sexual traits with the onset of SSD suggests that endocrine regulation has a role, potentially involving sex-specific hormonal shifts. Yet, hormonal control of growth in spiders is still poorly understood. Trabalon & Blais (2012) reported a spike in ecdysteroids in males just before maturity, which may contribute to growth arrest, whereas females might sustain or upregulate the expression of growth-related genes. Such putative shifts could help explain the emergence of dimorphism through sex-specific regulation of growth. While our results suggest that SSD emerges when males reach the subadult stage, Cordellier et al. (2020) propose that the developmental basis of sex differences must be initiated earlier, prior to the pre-subadult stage, implying that the physiological groundwork for SSD might have been laid well before it becomes morphologically visible. Future studies tracking gene expression across early instars may identify the molecular and regulatory cues that initiate sex-specific divergence in growth and morphology.

Family effects on growth were pronounced and strongly sex-specific. Family-specific trajectories indicated that the onset of female growth acceleration and male growth deceleration varied widely among families, occurring as early as ∼70 days in some and as late as ∼110 days in others. This substantial temporal variation implies that the timing and magnitude of sex-specific growth divergence are strongly family-dependent. Consistent with this notion, family-level analyses revealed marked sex differences in the correlation between initial size and subsequent growth. In females, families with larger-than-average size at the reference age tended to show slower subsequent growth, whereas families with smaller initial size tended to grow more rapidly. In males, by contrast, this correlation was weak and statistically indistinguishable from zero, suggesting that such compensatory growth patterns are largely absent or much less consistent in the male sex. These complex, family-dependent patterns are likely to complicate mechanistic studies of the developmental basis of SSD. The results suggest that, first, comparisons between sexes should be conducted within families to enhance statistical power, and, second, several families per species should be studied to achieve better generalization at the species level.

### Growth and seasonality

Hatching date significantly influenced carapace width at the average developmental age in both sexes. Individuals hatching during increasing day length attained larger carapace widths, followed by a decline and later rise, indicating that seasonal cues shape growth trajectories in *N. cruentata*. Although the study population has overlapping generations and is not kept under strictly natural seasonal environmental conditions, variation in climate and food availability persists, and growth appears to follow day length predictively. Hatching during increasing day length may permit prolonged growth and larger size when days are longer, whereas hatching during decreasing day length may constrain growth and reduce size, mitigating the ’end-of-season penalty.’ These patterns suggest predictive developmental plasticity, allowing individuals to adjust growth according to seasonal constraints, and align with previous reports of seasonal variation in body mass in *N. cruentata* (Kralj-Fišer et al. 2023) and other species (Higgins 2000; Chown & Klok 2003; Lissowsky et al. 2021). Interestingly, the seasonality in nature could be correlated with prey abundance, whereas in the laboratory the prey abundance remains equal across seasons. This suggests that seasonality in growth patterns is not directly regulated by prey abundance and food intake, but rather by seasonal cues, such as photoperiod.

Our multivariate analyses show that carapace width is an excellent proxy for body mass in *N. cruentata*, with very high family-level, residual, and phenotypic correlations in both sexes. Consequently, the pronounced seasonal patterns we document for carapace width closely reflect seasonal variation in body mass/condition, thereby directly linking our conclusions to previous work on seasonal dynamics of the easier-to-measure body mass in this species.

### Conclusions

Our study demonstrates that extreme SSD in *N. cruentata* originates from females sustaining growth long after male growth ceases, with divergence becoming apparent at the male subadult stage. This late divergence implicates emerging hormonal and/or molecular regulation, rather than early juvenile differences, as proximate drivers of SSD. Seasonal variation further modulates final size, highlighting the interplay between intrinsic developmental programs and environmental cues that should be accounted for in studies on SSD.

By identifying when male and female growth trajectories diverge, we provide a framework for future research into the physiological and genetic mechanisms underlying SSD. The complex patterns between sexes have a strong family dependence that stresses the importance of explicitly accounting for family structure in research on SSD.

## Acknowledgments

We thank Ž. Vehovar for helping with spider and data recording.

## AUTHOR CONTRIBUTIONS

**Conceptualization**– formulating the research goals and aims: Simona Kralj-Fišer (SKF), Paul Vincent Debes (PVD).

**Methodology**– designing methods, models, or experiments: SKF

**Investigation / Data collection**– performing the experiments, fieldwork, or lab work: Rok Golobinek (RG)

**Formal analysis**– statistical or computational analysis of data: PVD

**Resources**– provision of study materials, specimens, or other analysis tools: SKF, RG

**Data curation**– managing, cleaning, and maintaining data for reuse: Tim Prezelj (TP), PVD

**Writing – original draft**– preparing and writing the initial manuscript: SKF, TP, PVD

**Writing – review & editing**– critical review, commentary, and revision: SKF, PVD

**Visualization**– preparing figures, tables, data presentation: SKF, PVD

**Supervision**– oversight of the research project: SKF, PVD

**Project administration**– coordination, management: SKF

**Funding acquisition**– securing financial support for the project: SKF

## Supplementary material

**Table S1:** Number of times female and male spider individuals were measured for carapace width and the total number of measurements per sex and life stage.

| Sex | 1x | 2x | 3x | 4x | juvenile | subadult | adult |
| --- | --- | --- | --- | --- | --- | --- | --- |
| Female | 119 | 65 | 72 | 2 | 161 | 28 | 207 |
| Male | 145 | 500 | 13 | 0 | 117 | 497 | 566 |

